# Variomes: a high recall search engine to support the curation of genomic variants

**DOI:** 10.1101/2021.05.29.446224

**Authors:** Emilie Pasche, Anaïs Mottaz, Déborah Caucheteur, Julien Gobeill, Pierre-André Michel, Patrick Ruch

## Abstract

Precision oncology relies on the use of treatments targeting specific genetic variants. However, identifying clinically actionable variants as well as relevant information likely to be used to treat a patient with a given cancer is a labor-intensive task, which includes searching the literature for a large set of variants. The lack of universally adopted standard nomenclature for variants requires the development of variant-specific literature search engines. We develop a system to perform triage of publications relevant to support an evidence-based decision. Together with providing a ranked list of articles for a given variant, the system is also able to prioritize variants, as found in a Variant Calling Format, assuming that the clinical actionability of a genetic variant is correlated with the volume of literature published about the variant. Our system searches within three pre-annotated document collections: MEDLINE abstracts, PubMed Central full-text articles and ClinicalTrials.gov clinical trials. A variant synonym generator is used to increase the comprehensiveness of the set of retrieved documents. We then apply different strategies to rank the publications. We assess the search effectiveness of the system using different experimental settings. Experimental setting 1: The literature retrieval task is tuned and evaluated using the TREC Precision Medicine 2018 and 2019 benchmarks consisting respectively in 50 and 40 topics. Almost two thirds (62%) of the publications returned in the top-5 are relevant for clinical decision-support. Experimental setting 2: The evaluation of the variant prioritization task is based on a manually-created benchmark composed of eight patients for a total of 756 variants. For each patient, we used both their complete set of variants and tumor board reports. Our approach enabled identifying 81.8% of clinically actionable variants in the top-3. Experimental setting 3: A comparison of Variomes with LitVar, a well-known search engine for genetic variants is performed. Variomes was able to retrieve on average 90.8% of the content, while LitVar retrieved on average 58.6%. Out of the 9.2% articles, which are “missed” by Variomes, a per error analysis suggests that they are artefacts. To conclude, we are proposing here a competitive system to facilitate the curation of variants for personalized medicine.

## 1. Introduction

Advances in personalized medicine make it now possible to select a treatment targeting specific tumor variants. Indeed, based on the tumor’s molecular profile coupled with clinical information such as the diagnosis, it is possible to better determine which treatment resulting in a likely favorable response can be proposed. To this extent, a tissue sample is sequenced, resulting in the identification of hundreds of variants. The clinical experts, assisted by bioinformatics tools, are then in charge of determining which variants are actionable, i.e. are likely to result in a better or worse prognosis and derived treatment response. It results in the generation of a tumor board report which can be used by physicians for the treatment of the patient.

The identification and interpretation of clinically actionable variants is a critical bottleneck. Indeed, it is necessary to look for evidence in genomic variant knowledgebases such as oncoKB, COSMIC, CIViC, as well as in the literature. These high-quality resources rely on manual curation. However, since manual curation is not scalable, information on curated variants is sometimes incomplete or out-of-date. Moreover, their coverage is very diverse. Lee *et al*. [1] reported that only eight variants overlapped over six well-known cancer genomic variant databases. Scientific literature is thus an indispensable source of content. However, screening out scholarly publications poses a number of challenges. Curators must cope with large and increasing volumes of publications: Lee *et al*. [1] showed that among the last five years 50,000 publications containing genomic variants were published every year. Moreover, the information is “hidden” in unstructured text in which the genetic variants, but also the other relevant entities (e.g. genes, diseases, demographic information, etc.), are labelled in diverse forms such as synonyms and abbreviations. While a few variants received massive attention, most of them - the so-called Variant or Unknown/Uncertain Significance (VUS) - are described in a handful of published reports. For instance, only 13 MEDLINE abstracts mention the genomic variant P871L of the Breast Cancer 1 gene and gene product. However there are 37 additional abstracts of potential interest in which the variant is named with alternative forms, such as p.Pro871Leu or c.2612C>T. By using PubMed-like search engines, it is therefore necessary to multiply the queries to avoid missing an important publication. In addition, when the number of publications is large (e.g. for highly studied variants), the triage of the literature to select the most relevant documents is a tedious task.

In recent years, search engines dedicated to variant-related tasks have aroused the interest of the Natural Language Processing (NLP) community. In particular NLP competitions have accelerated the investigation of more sophisticated approaches. Since 2017, TREC Precision Medicine track [2,3] has been proposing a task aiming at finding relevant publications given a particular case containing a genetic variant, a disease and demographic information. The best resulting systems were able to retrieve almost two thirds of relevant publications in their top-10 results.

When developing a search engine to retrieve literature for genomic variants, four aspects should be taken into account: the literature collections, the normalization of variant names, the type of search algorithms and the literature triage. Regarding the literature collections, MEDLINE proposes a corpus of more than 30 million citations, out of which about 800,000 are related to mutations [4]. While abstracts can be considered as sufficient for variant prioritization, full-text access is needed to support the clinical interpretation of variants. Indeed, information related to genomic variants are often mentioned in the body of the scientific reports, including results and tables. According to Jimeno Yepes and Verspoor [5] a significant subset of the information related to genomic variants is even reported in supplementary data. The normalization of variants’ names into a single form is an essential step. Indeed, variants can be represented in a multitude of standard formats (e.g. amino acids can be represented using one-letter codes or three-letters codes) or even by using non-standard expressions as described by Yip *et al*. [6]. Lee *et al*. [1] reported that 76% of the variants mentioned in the literature do not follow the standard HGVS nomenclature. Thus, the use of variant-specific name entity recognition (NER) tools is essential.

Various variant-specific search engines have been developed in recent years. We present here a brief overview. LitVar [7], one of the most cited ones, uses both abstracts and full-texts. Using information matching, it returns a chronologically-ordered set of publications. variant2literature [8] not only uses full-text articles but also processes supplementary data, in particular tables represented as images in PDF. Like LitVar it returns literature in a chronological order. Overcoming this chronological ordering limitation, VIST [9] uses a support vector machine model to rank publications by relevance. However, this tool does not cope with full-text articles: only abstracts are used. Nevertheless, it is one of the rare tools including searching for clinical trials. LitVar, variant2literature and VIST all rely on the use of tmVar [10] for the recognition and normalization of variant names in both publications and users’ queries. Other types of approaches focus on the literature triage, such as LitGen [11], which collects publications returned by LitVar and filters them by types of evidence. Further, Lv *et al*. [12] proposes one of the only methods not based on tmVar or LitVar. It is based on a Knowledge-enhanced Multi-channel CNN model to identify relevant publications both in abstracts and full-texts.

In this context, we designed Variomes [13], an original application to support the search of human variants. The system can be used as a literature triage system in the same way as LitVar. It can also be used to prioritize variants (e.g. from a Variant Calling File) to facilitate the identification of clinically actionable variants. When the system is used to rank variants, it assumes that the clinical actionability of a variant is directly correlated with the volume of literature published about the variant. The ranking of variants consists of establishing a score for each variant by summing the scores - more precisely the Retrieval Status Value as returned by the search engine, see [14] for more information - of publications retrieved for each variant. On the contrary, a variant without clinical significance (e.g. silent mutation) will result in no or very few citations.

Our tool aims to facilitate the annotation of the variants by curators by suggesting them a set of publications of interest. Variomes enables searching the biomedical literature. The collections are pre-processed with a set of medical terminologies [15]. At query time, user queries are automatically processed to map keywords to the terminologies and expand genetic variants using a dedicated variant expansion system [13]. Finally, different strategies are investigated to maximize the performance of the literature triage based on the tuning of an optimal ranking function [16,17]. Variomes is available through a user-friendly interface as well as through a set of APIs. Finally, it is also integrated within the SVIP curation platform [18], a national Swiss repository for clinically verified variant annotations in oncology.

## 2. Data and methods

Our approach is based on the use of three collections of scientific literature: abstracts from MEDLINE, full-text articles from PubMed Central, and clinical trials from ClinicalTrials.gov.

### 2.1 System’s architecture

The collection is first normalized with a set of terminologies in order to ease the matching of user’s information requests. The following terminologies were selected: neXtProt [19] for genes, NCI Thesaurus [20] for diseases and DrugBank [21] for drugs. Pre-processing the collections enables first to retrieve results faster thanks to pre-computed indexes and second to increase the recall. Indeed, querying the collections through the annotations permits retrieving not only the exact term, but also its synonyms as well as string variations. For instance, while querying the MEDLINE collection using the keyword *BRAF* returns 15,907 hits, querying the annotations using *NX_P15056* (i.e. the unique neXtProt identifier corresponding to the *BRAF* gene) returns 17,191 documents, thus increasing the recall by almost +8%. Indeed, the annotations recognized not only *BRAF* and its official synonyms *BRAF1* and *RAFB1*, but also syntactic variations such as *B-RAF*.

Documents and annotations are loaded into a MongoDB document database and indexed into an ElasticSearch Index. Our system uses the ElasticSearch index for querying, while the MongoDB database serves for annotations-based re-ranking as well as for enriching the display of the documents.

The search engine is based on a two-steps system. The first step focuses on recall as it aims at gathering a large set of documents related to a particular case. The second step focuses on precision as it attempts to properly rank the set of documents.

Each variant query can be represented as a triplet: a variant in a gene for a given diagnosis. To collect the most comprehensive set of abstracts, an Elasticsearch query is generated. This query is composed of three “must” clauses (i.e. one clause for each entity of the triplet) that must appear in the matching documents. For the disease and the gene clauses, two “should” sub-clauses are defined and at least one of the clauses must appear in the matching documents: the exact query term is searched in the publications text and the corresponding unique identifier is searched in the annotations. For the variant clause, a set of “should” sub-clauses are generated: one for the exact query term and one for each of its synonyms generated by the SynVar service. SynVar is a synonym generator for single nucleotide polymorphisms (SNP) [13]. It provides descriptions of the variant at other levels (e.g. genomic level), as well as syntactic variations encountered in the literature. It proposes up to 50 synonyms for a variant, with descriptors at the protein, transcript and genomic levels.

However, the triplet-based query is sometimes too specific and documents not strictly targeting a given triplet might still be valuable, so that a constraint relaxing strategy is needed. For instance, a document about the given variant in the given gene but for another diagnosis - e.g. melanoma instead of breast cancer - may still be valuable from a clinical point of view. Moreover, while full-text articles reporting on treatments usually mention all the information regarding the disease, gene and variant, it is not always the case with abstracts. Thus, our system also collects documents with decreasing levels of specificity. Three additional queries are generated, each omitting one of the entities of the triplet. The respective outputs of these additional queries are linearly combined. Weights attributed to each additional query’s output are automatically tuned using Fox and Shaw’s methods [22].

The re-ranking of publications is based on a matrix of scores. A score is calculated for each ranking strategy: 1) the relevance score provided by ElasticSearch, 2) a score based on the constraint relaxing strategy described above, 3) a score based on the density of some specific named-entities in the document, 4) a score based on the demographic concordance if demographic information is available in the query and 5) a score based on the density of some predefined keywords. These scores are then combined with different weights to generate a final score for each publication. Finally, the scores of documents published in languages other than English are downgraded to be returned after English documents.

The score for the density of some specific named-entities is based on the number of occurrences of gene descriptors, disease descriptors and drug descriptors. To accelerate the scoring of the retrieved documents at query time, the collection is pre-annotated with a large set of named-entity types. All weights are determined by direct search using TREC benchmarks. The score for the demographic concordance is based on age-groups and genders extracted from the MeSH terms associated with each MEDLINE record. Currently, this approach is applied only on the MEDLINE collection. When an abstract is targeting the required gender and/or age-group, a positive boost is applied. The score based on predefined keywords aims to classify a document as being related to precision medicine or not. A list of positive stemmed keywords (e.g. *treat*) and negative stemmed keywords (e.g. *marker*), has been manually defined through a manual screening of a subset of documents. The presence of positive keywords in a document will improve the scoring, while the negative keywords occurrences will decrease its relevance.

### 2.2 Experimental evaluation setting

We perform three types of evaluation of the services: 1) as a literature triage system to support the curation of a given variant using different standard metrics, which balance both recall and precision; 2) as a variant triage system using a VCF as input with focus on precision; 3) a comparison between our system and LitVar with focus on recall.

In the absence of benchmarks with VCF queries, the system is initially tuned to perform a literature triage task; it means that most queries contain only a single variant. A benchmark provided by the TREC Precision Medicine track is used: the TREC PM 2018 benchmark [2] for the tuning and the TREC PM 2019 [3] benchmark for the evaluation. Both benchmarks consist of semi-structured synthetic cases created by precision oncologists at the University of Texas MD Anderson Cancer Center. Each topic mentions a disease, one or several mutated gene(s) and, optionally, some demographic information. The tuning and evaluation benchmarks contain respectively 50 and 40 topics. In addition, we also perform an experiment to assess the use of synonyms of variants for literature triage. To this extent three runs are performed: querying the system with all the variant synonyms, including gene to protein translation (default settings as described above), querying the system with strict protein synonyms (i.e. *V600E* is expanded to *Val600Glu* but not to *1799T>A*) and finally querying the system with no synonym.

Further, the variant prioritization is evaluated using a dataset of 756 variants, originating from eight patients, a partial set of the SwissMTB study [23]. For each patient, called variants and manually curated tumor board reports are evaluated. A topic consists of a diagnosis, a set of variants containing on average 94.5 (16 - 439) single nucleotide variants (SNV) and optionally a gender and/or an age. For each topic, the VCF has been pre-processed to generate a set of gene and variant pairs. Only single nucleotides at the protein level are selected. The tumor board reports are used to define the variants which should be top-ranked by the system. Five topics contain one relevant SNV and three topics contain two relevant SNVs.

Furthermore, a comparison between LitVar and Variomes is also performed to evaluate the recall of our system. For this task, we did not use the constraint relaxing strategy of our system. A set of 803 queries containing variants in BRCA1 and BRCA2, originating from BRCAExchange [24] is used. Queries are sent to both systems and an automatic comparison of the returned documents is performed. In addition to the quantitative comparison, we also perform a qualitative analysis on a random subset of queries. The manual analysis of the results aims to identify what features could explain such differences.

## 3. Results and discussion

### 3.1 Tuning of the system

The tuning of the system is based on five steps: 1) the scoring of the constraint relaxing strategy, 2) the scoring of the density of named-entity types, 3) the scoring of the demographic concordance and 4) the scoring of the predefined keywords and 5) the linear combination of the four strategies. We present here the best settings selected for each step. The best settings are selected based on the maximization of the R-Prec (R-Precision), the P5 (Precision at rank 5) and the infNCDG (inferred non discounted cumulative gain). R-Prec returns the number of relevant documents returned in the top-R documents, where R corresponds to the number of relevant documents for the query. P5 represents the proportion of relevant documents retrieved in the top-five results. Finally, infNDCG reflects the gain brought by a document based on its position in the ranked results.

The constraint relaxing strategy was tested by attributing a weight between 0.0 and 1.0 to each of the three relaxed queries. The best results were obtained when using a weight of 0.95 for the query containing the disease and the gene, a weight of 0.07 for the query containing the disease and the variant and a weight of 0.05 for the query containing the gene and the variant.

The scoring of the density of named-entity types was tested by attributing a weight between 0.0 to 1.0 to the following named-entity types: gene, disease and drug. The best results were obtained when using a weight 0.97 for the disease, 0.51 for the gene and 0.57 for the drug.

The scoring of the demographic concordance was tested by attributing a weight to the age score and the gender score, between 0.0 and 1.0. The age and gender scores are calculated by attributing a strong bonus to documents matching the requested age and gender (bonus between 0.7 and 1.0) and a moderate bonus to documents not discussing the age or gender (bonus between 0.1 and 0.4). The best results were obtained when using a weight of 0.7 for the age and 0.5 for the gender, as well as a bonus of 0.7 for documents matching the age, 0.4 for documents not mentioning the age, a bonus of 0.7 for documents matching the gender and 0.4 for documents not mentioning the gender.

The scoring of the predefined keywords consisted first to define a list of positive and negative stemmed keywords. Two lists were tested for each modality: *treat;drug;therap;prognos;surviv* and *treat;drug;therap* for positive keywords; *immuno;marker;detect;sequencing* and *immuno;marker;detec*t for negative keywords. A bonus between 0.0 and 1.0 was attributed to each occurrence of a positive keyword, while a penalty between 0.0 and −1.0 was attributed to each occurrence of a negative keyword. The best results were obtained when using the following keywords lists for positive and negative keywords: *treat;drug;therap;prognos;surviv* and *immuno;marker;detect*. The best settings occurred when using a bonus of 0.2 for the positive keywords and a penalty of −0.1 for the negative keywords.

Finally, the weight attributed to each of these strategies was defined by testing weights between 0.0 and 1.0. The ElasticSearch score was granted a weight of 1.0. The best results were obtained when giving a weight of 0.65 to the constraint relaxing, 0.1 to the named-entity types density, 0.05 to the demographic concordance and 0.1 to the predefined keywords.

### 3.2 Literature triage

The settings defined in the previous section were used to evaluate the literature triage task. The system resulted in a R-Prec of 32.5%, an infNDCG of 49.8% and a P5 of 62%, which means that almost two thirds of the top-5 returned abstracts are judged relevant. One third of the relevant documents are retrieved in the top-R documents.

An analysis of the P5 topic per topic is presented in Figure 1. The system performs the best with queries related to SNV. Indeed, 80% of the SNV queries resulted in a P5 equal or greater to 80%. However, queries related to gene fusion (classified as others in the figure below) do not perform well, with a P5 between 20% and 60%.

**Figure 1:**
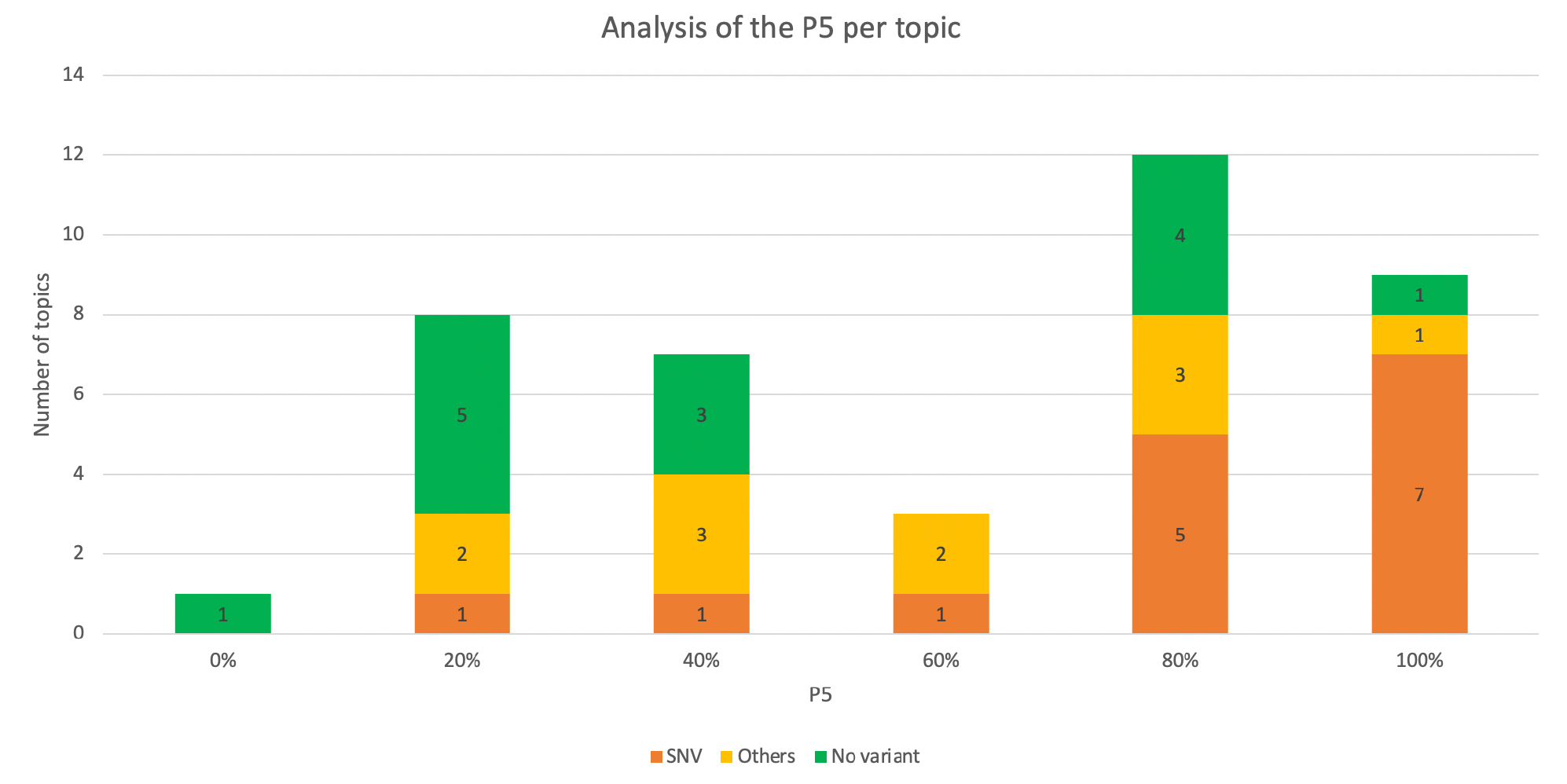
Distribution of topics (number and type) according to the P5 values.

Our system is composed of two steps: collecting a set of abstracts and re-ranking these abstracts. While the second step is important to prioritize the literature, the first step is mandatory: the relevant abstracts must be broadly captured in our set to be properly ranked. We thus investigate which proportions of relevant abstracts have been successfully retrieved by our system, at whatever ranks they were returned. However, for queries resulting in a large amount of documents, only the top-1000 documents are retrieved. For more than half of the topics (22/40), at least 70% of the relevant documents are retrieved in our abstracts set. For such topics, the focus should be put on improving the ranking. However, for twelve topics, less than half of the relevant documents are retrieved. Further investigations are needed to analyze such abstracts and define possible actions to improve their gathering, such as improving the annotations of genes or diseases, expanding the diseases to parent and/or children diseases, defining better synonym lists for genetic variants, etc.

The results of the three runs to assess the effect of variant’s synonyms are presented in Table 1. We observed that using all the synonyms proposed by the SynVar service surprisingly decreased the precision by −2.3%. Indeed, we obtained a P5 of 63.5% when no synonyms were used to query ElasticSearch. However, using only the basic synonyms (i.e. those obtained without any mapping) resulted in the best P5 (64%). It appeared from discussions with the TREC PM organizers that complex synonyms of variants were not taken into account by the assessors. This is a limitation of the TREC benchmark since publications retrieved using complex synonyms are of equal clinical significance for the curation of variants.

**Table 1:** comparison of the effect of variants synonyms on the results of literature triage

|  | **P5** | **R-Prec** | **infNDCG** |
| --- | --- | --- | --- |
| **Run with all variants synonyms** | 62% | 32.5% | 49.8% |
| **Run with no variants synonym** | 63.5% | 32.4% | 49.6% |
| **Run with basic variants synonyms** | 64% | 32.7% | 50.1% |

### 3.3 Variants prioritization

The variant prioritization resulted in a P5 of 25%, which means that a quarter of the variants returned in the top-five are judged as clinically actionable according to the tumor board. It is to be noted that the other variants returned in the top-five are not necessarily irrelevant but have not been assessed by the tumor board. Because the relevance judgments only contain one or two relevant variants per topic, the R-Prec is a more appropriate metric: 71.4% of the variants returned in the top-R results are relevant. Out of the eleven variants reported in the tumor board, 81.8% of the variants were returned in the top-3: five were returned at rank #1, three at rank #2 and one at rank #3. For two variants, no literature was found; thus, suggesting that recall remains the main challenge for such a task.

### 3.4 Comparison with LitVar

Quantitative results of the comparison of LitVar and Variomes are presented in Table 2. 7110 documents were retrieved in total with 3701 documents (52%) in common between the two systems.

**Table 2:** quantitative results of the comparison of Variomes and LitVar

|  | **Variomes** | **LitVar** | **Union** |
| --- | --- | --- | --- |
| **Average number of documents retrieved per query** | 7.4 | 6.1 | 8.9 |
| **Average percentage of the content retrieved per query** | 90.8% | 58.6% | 100% |
| **Total number of documents retrieved** | 5915 | 4896 | 7110 |
| **Number of queries with no results** | 144 | 253 | 259 |
| **Number of queries with more results than the other systems** | 462 | 80 | - |

Variomes was able to retrieve on average 90.8% of the published content (i.e. results retrieved by LitVar + results retrieved by Variomes), while LitVar retrieved on average 58.6%. For 462 queries (57.5%), Variomes retrieved more documents than LitVar, while for 80 queries (10%), LitVar returned more documents. 140 queries (17.4%) returned no results whatever systems are used, which again suggests that recall remains a challenge for variant retrieval. In addition to these 140 queries, LitVar and Variomes also returned no document for respectively 113 and 4 queries.

Ten randomly selected queries were manually analyzed (Table 3) in an attempt to better understand where lies the respective power of Variomes and LitVar. For two queries, only Variomes returned results. For five queries, Variomes returned the total set retrieved by LitVar, as well as one to eleven additional documents. For the other three queries, Variomes missed one to three documents retrieved by LitVar, but for two of these queries, it also proposed one to three documents not retrieved by LitVar. In total, 68 documents were retrieved for the five queries, with 34 documents in common among both systems.

**Table 3:** ten randomly selected queries for the manual comparison of Variomes and LitVar.

|  | Results by Variomes | Results by LitVar | Total results | Only in Variomes | Only in LitVar |
| --- | --- | --- | --- | --- | --- |
| BRCA1 (L246V) | 10 | 8 | 10 | 2 | 0 |
| BRCA1 (I1044V) | 2 | 1 | 2 | 1 | 0 |
| BRCA2 (V3091I) | 4 | 2 | 4 | 2 | 0 |
| BRCA2 (G2813E) | 3 | 1 | 3 | 2 | 0 |
| BRCA1 (S1497A) | 5 | 0 | 5 | 5 | 0 |
| BRCA1 (*R3052W*) | 27 | 16 | 27 | 11 | 0 |
| BRCA2 (R2842L) | 3 | 5 | 6 | 1 | 3 |
| BRCA1 (D67E) | 5 | 4 | 7 | 3 | 2 |
| BRCA1 (A1830T) | 2 | 3 | 3 | 0 | 1 |
| BRCA2 (Q3066E) | 1 | 0 | 1 | 1 | 0 |

Variomes failed to retrieve seven documents, which were retrieved by LitVar. In three of these documents, the variant is observed in the reference section (i.e. in the title of another publication). In such cases, the relevance of the citing article is doubtful. In the four other documents, the variant highlighted by LitVar is not the one mentioned in the query (e.g. *R2842H* instead of *R2842L*). Indeed, LitVar is using RSIDs, which combine variants occurring at the same locus. This is also the main cause observed for the two queries where Variomes retrieved fewer articles than LitVar (*BRCA1 R1443G* and *BRCA1 A1708V*). Thus, we can consider from this error analysis that no or very few relevant articles were really missed by our system compared to LitVar. The fraction of articles not found by Variomes (9.2%) is in fact an experimental artefact; thus, it is estimated that Variomes’ recall is about 70% higher than LitVar.

To evaluate the quality of documents retrieved solely by Variomes, we merged the 28 pairs of variants/documents from Table 3 together with 50 other randomly selected documents. We reached a precision of 86%. Regarding the relevant documents specific to Variomes, in most cases, the variant was present, but with a different form (e.g. *9154C>T→G* for *R3052W*). We can thus deduct that the variant expansion services of Variomes (so-called SynVar and whose API can be accessed independently from Variomes) service proposes a larger set of patterns for variant synonyms. Further, some of the retrieved documents were functional assays of human BRCA genes performed in various cells such as mouse stem cells, bacteria, yeast and human cancer cells. They may not have been considered as “human” research by LitVar while they could clearly be of interest for some clinical research. Regarding the non-relevant documents, they had two origins. First, some variants were in reference to another gene. Second, some documents were retrieved through a mentioned RSID but were about another variant at the same position.

### 5.5 Availability

The Variomes service is publicly available at http://candy.hesge.ch/Variomes/. The user can either query the service with a single variant (or a combination of several variants) or he/she can upload a file containing a list of variants. A few parameters enable to personalize the search: specification of the timeline, addition of keywords to re-rank the documents, specification of facultative entities (e.g. disease), etc. The variants are then returned in a ranked table. The user can select a variant to access the retrieved literature (MEDLINE abstracts, EuropePMC full-texts and clinical trials). Each document is displayed with highlighted annotations, as shown in Figure 2. The user can mark publications of interest. Thus he/she can at the end generate and export a report (i.e. JSON or HTML) that summarize all the variants of interest with publications selected as relevant by the user. In addition, public APIs are also available for the major functionalities: e.g. retrieving ranked literature for a given variant, retrieving annotated literature and retrieving variant synonyms. APIs are described here: http://candy.hesge.ch/Variomes/api.html. Finally, Variomes services are also integrated with the SVIP prototype.

**Figure 2:**
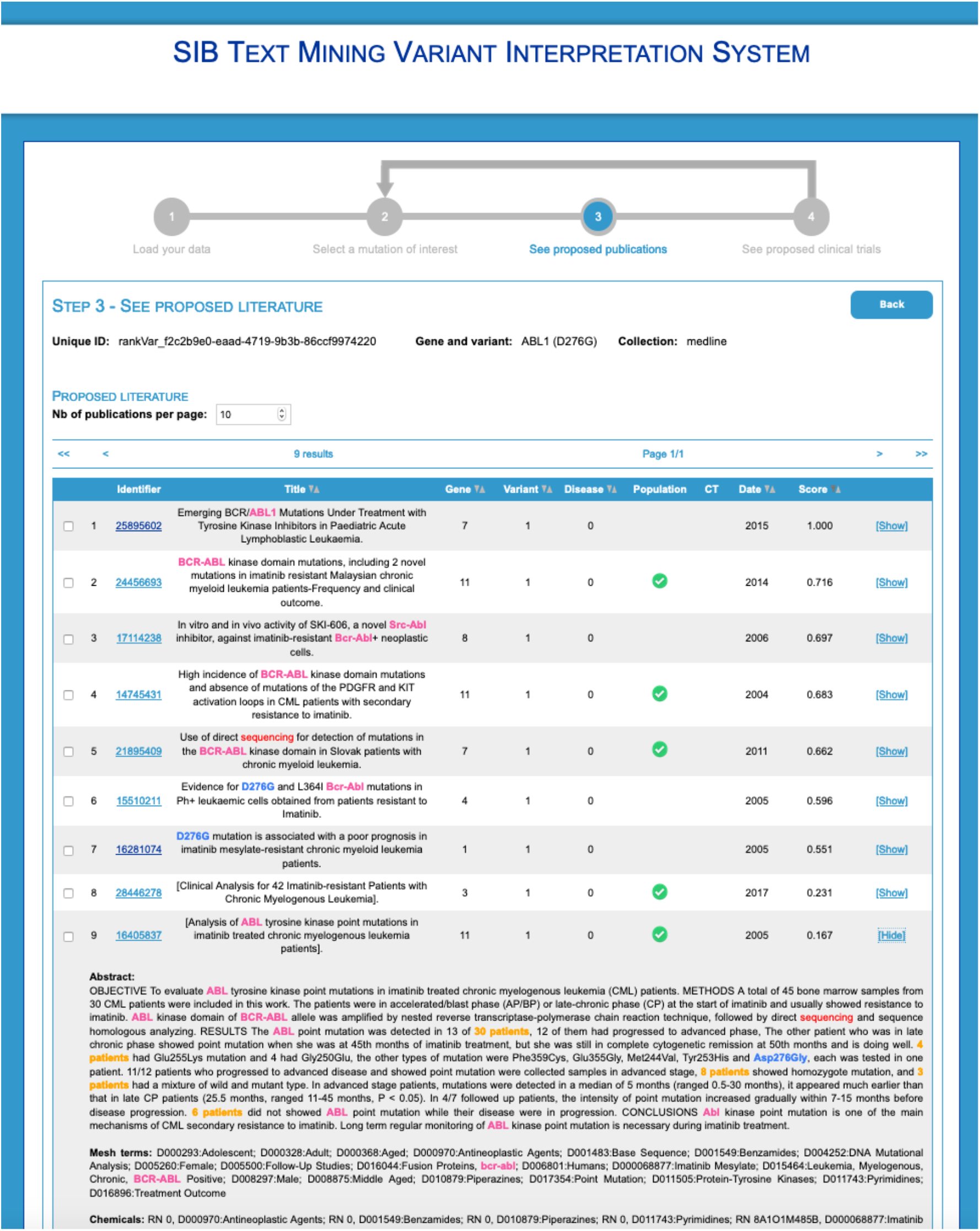
Example of the Variomes demonstrator for the abstracts retrieved for the query ABL1 D276G.

## Conclusion

We are proposing an efficient tool for retrieving literature associated with variants. The system defines a new state of the art for retrieving genomic variants. For literature triage, our system was able to retrieve almost two thirds of the relevant publications in the top-five. The Variomes system is particularly efficient with single nucleotide variants, where the P5 was greater than 80% for most of the SNVs queries. Such a result is consistent with the targeted application of the tool: SNV accounts for the vast majority of contents in VCF files, whether they are based on gene panels or on WGS/WES. We are now including other types of variants in the system to better cover the needs of personalized medicine. The expansion of the tool beyond somatic mutations is also a work area. Germline mutations, as well as mutations in non-human genomes, including viruses, are currently being considered.

In comparison to LitVar, one of the most popular variant-specific literature search tools, we obtained competitive results. Indeed, our system retrieved more relevant documents than LitVar. On the one hand, thanks to the various patterns proposed by the SynVar service, more relevant documents were retrieved. On the other hand, our system offers a larger collection as it covers not only human research but also animal studies. In addition, our system is also more focused on the exact variant requested by the curator, while LitVar returns any variation occurring at the same locus, which may result in an overload of publications to curate. Further, the literature content of Variomes, which is powered by the SIB Literature Services (SIBiLS) [15], is also wider than LitVar as it provides access to complementary data sources, such as the ClinicalTrials.gov, which are directly relevant for variant curation, functional assays of human genes in non-human cells, or collections describing biodiversity-related variants, from mammals (e.g. Bats, Pangolin, etc.) to viruses, but which could have relevance for healthcare. Such additional content was not used in the present study but could potentially broaden the scope of the variant search.

As future work we consider three aspects. First, we would like to investigate the use with supplementary data. Indeed, as shown by Jimeno Yepes and Verspoor [5], some variants only appear in such data, which may explain the silence of about 20% of the queries. Second, identifying textual evidence in publication to re-rank publications might have a positive impact on the literature triage [25, 26]. Finally, pre-trained language and ensemble learning models [27] could be opportunely used to provide the curator with a more focused evidence passage to support the curation work of mutation databases [18]

To conclude, the system we developed has the potential to significantly propel variant curation. It is however to be noted that such a system is neither intended to replace human curators, nor clinical expertise, but rather to support these professionals by cutting down the cost of the manual triage of the literature.

## Acknowledgments

This Swiss Variant Interpretation Platform (SVIP) project has been supported by the Swiss Personalized Health Network (SPHN) and the BioMedIT infrastructure, see https://svip.ch/. SVIP uses an integrated version of the Variomes services to support the curators of the clinical database. We would like to thank the SVIP project team members of the following groups: Clinical Bioinformatics Unit of NEXUS Personalized Health Technologies, SIB Clinical Bioinformatics and Swiss-Prot group, namely Daniel Stekhoven, Valérie Barbié, Anne Estreicher, Livia Famiglietti, Faisal Al Quaddoomi, David Meyer, Linda Grob, Franziska Singer and Nora Toussaint. The work presented in this report is built on top of SIBiLS, the SIB Literature Services, which is supported by the Elixir Data Platform. This work also benefited from discussions with Melissa Cline.

